# Symmetric assembly and disassembly processes in an ecological network

**DOI:** 10.1101/042572

**Authors:** Jason M. Tylianakis, Laura B. Martínez-García, Sarah J. Richardson, Duane A. Peltzer, Ian A. Dickie

**Affiliations:** Centre for Integrative Ecology, School of Biological Sciences, University of Canterbury, Private Bag 4800, Christchurch 8140, New Zealand.; Department of Life Sciences, Imperial College London, Silwood Park Campus, Buckhurst Road, Ascot, Berkshire SL5 7PY, United Kingdom; Bio-protection Research Centre, School of Biological Sciences, University of Canterbury, Private Bag 4800, Christchurch 8140, New Zealand.; Landcare Research, PO Box 69040, Lincoln 7640, New Zealand; Department of Soil Quality, Wageningen University. P.O. Box 47. Wageningen 6700 AA, The Netherlands

**Keywords:** community assembly, succession, ecosystem development, retrogression, mutualist network, mycorrhizal symbiosis, preferential attachment

## Abstract

The processes whereby ecological networks emerge, persist and decay throughout ecosystem development are largely unknown. Here we study networks of plant and arbuscular mycorrhizal fungal (AMF) communities along a 120,000 yr soil chronosequence, as they undergo assembly (progression) and then disassembly (retrogression). We found that network assembly and disassembly were symmetrical, self-reinforcing processes that together were capable of generating key attributes of network architecture. Plant and AMF species that had short indirect paths to others in the community (i.e. high centrality), rather than many direct interaction partners (i.e. high degree), were best able to attract new interaction partners and in the case of AMF species, also to retain existing interactions with plants during retrogression. We then show using simulations that these non-random patterns of attachment and detachment promote nestedness of the network. These results have implications for predicting extinction sequences, identifying focal points for invasions, and suggesting trajectories for restoration.

## Introduction

The arrangement of species interactions in ecological networks can determine the stability and functioning of ecosystems (Bastolla *et al*. 2009; Thebault & Fontaine 2010; Thompson *et al*. 2012). However, the processes that generate and maintain network structure during ecosystem development are unresolved. The generally accepted hypothesis for complex network assembly is that new nodes (e.g., species) preferentially connect with those that are linked to many immediate neighbors (Barabási & Albert 1999) (i.e. generalist nodes with high ‘degree’). This ‘preferential attachment’ was predicted (Jordano *et al*. 2003), then shown empirically (Olesen *et al*. 2008), to occur during the assembly of pollination networks within a season. Moreover, preferential attachment could in theory promote nestedness, a network property that relates to network stability (Bastolla *et al*. 2009; Thebault & Fontaine 2010), such that networks would become more stable over time (Ponisio *et al*. 2017). In contrast, long-term changes to interaction networks have shown contradictory patterns, such that new interactions can primarily involve more-connected species in ant-plant networks (Díaz-Castelazo *et al*. 2010), less-connected species in pollination networks (Burkle *et al*. 2013), or simply be opportunistic based on available partners (Ponisio *et al*. 2017). Thus, the importance of preferential attachment based on species degree remains contentious for ecological networks.

Although studies on the generation of network architecture have largely focused on node attachment during network assembly, an opposing process of preferential detachment can theoretically produce the same architecture as preferential attachment (Salathé *et al*. 2005; König *et al*. 2012). This possibility is strengthened by empirical findings that pollination networks show preferential loss of less-connected species and their interactions during network decay (Aizen *et al*. 2012; Burkle *et al*. 2013). It is also possible that network architecture may be generated by preferential attachment during community assembly, and maintained by preferential detachment during disassembly. This would imply that the same rules may operate during both the assembly and disassembly of ecological networks (Bascompte & Stouffer 2009), rather than via different pathways, as would be expected if networks exhibit hysteresis (Suding & Hobbs 2009). To our knowledge, whether networks assemble and disassemble via the same mechanisms remains untested.

Here we study interaction network assembly and disassembly in a community of plants and arbuscular mycorrhizal fungi (AMF) (Montesinos-Navarro *et al*. 2012) throughout 120,000 years of ecosystem development. Arbuscular mycorrhizal fungi are essential mutualists of most terrestrial plants (Brundrett 2009), and have typical within-site diversity comparable to plant communities (Öpik *et al*. 2009). Moreover, the high importance of symbiotic interactions for determining AMF community composition (Martínez-García *et al*. 2015; Vályi *et al*. 2015) demands a focus on interactions (Powell *et al*. 2015) for these systems, within the context of soil nutrients which can also be important drivers of community structure (Krüger *et al*. 2015). First, we test whether network assembly and disassembly processes are random or related to species’ direct or indirect connections to others the network. Concomitantly, we test the hypothesis (Bascompte & Stouffer 2009) that the processes governing attachment during assembly mirror those for detachment during disassembly. Finally, we use simulations to determine how non-random assembly and disassembly processes impact upon key aspects of network architecture.

The first requirement for an interaction to take place is an encounter, and if co-occurrence rates between AMF propagules and plants were merely stochastic (Encinas-Viso *et al*. 2015), we would expect to see more frequent than random attachment to the most abundant partner species. In addition to such random encounter probabilities, species’ abundances may convey benefits to their interaction partners. For example, an established AMF species that has used its associations with many plant species to develop an extensive mycelium network could be an attractive partner for newly-arriving plants, because they would not need to pay the initial construction costs for the mycelium network but they would gain immediate access to a large soil volume (Newman 1988; Simard & Durall 2004). Likewise, high AMF diversity can reduce competition among plant species and provide insurance against variability in conditions (Moeller & Neubert 2016), such that plant species that interact with many AMF experience increased biomass (Wagg *et al*. 2011). This greater biomass could make AMF-generalist plants more attractive partners because they have the capacity to provide more carbon across a wider range of conditions. Selection for partner quality could therefore lead to traditional preferential attachment for abundant, generalist species (Barabási & Albert 1999; Olesen *et al*. 2008), and a mechanism for this selection could be post-infection selectivity by plants and AMF in their use of interaction partners (Kiers *et al*. 2011). In fact, preferential attachment to generalists could also occur if generality is simply an indicator of low partner selectivity.

The benefits of interacting with generalists may also depend in part on whether they share their interaction partners with other species. Although AMF typically interact with many plant species, there is growing evidence that their composition (Martínez-García *et al*. 2015) and fitness can differ across plant hosts (Ehinger *et al*. 2009). Host-plant selectivity may not necessarily be species specific, but rather groups of AMF can associate with ecological groups of plants (Davison *et al*. 2011). Closely-related AMF also tend to co-occur in a given location (Horn *et al*. 2014), and closely-related organisms tend to share interaction partners (Gómez *et al*. 2010). We therefore hypothesize that large interacting consortia of ecologically-similar, and in some cases phylogenetically related, plants and fungi will form at a given successional stage, with the dominant consortium potentially differing between early-successional generalist species and later-successional forest species (Öpik *et al*. 2009; Davison *et al*. 2011). Thus, at each stage of ecosystem development, newly-arriving fungi would have a higher probability of interacting with plants in the dominant ecological group at that stage, which will already interact with species from the dominant fungal group. These plant species would not necessarily be the most generalist individually, but they would share AMF partners with many other plant species in the dominant group at that stage (i.e. with the majority of species in the site). This high interaction-partner overlap with others in the network may also bring fitness benefits by reducing the distance that positive indirect effects must travel (Bastolla *et al*. 2009), which may make species with many short indirect paths to others more reliable interaction partners. We therefore hypothesise that the attractiveness of a species to new arrivals will depend on its distance (i.e. path length) from other species in the network, as a measure of the extent to which it could receive positive indirect effects by sharing partners with species from the dominant consortium (see Fig. S1 for our approach to measuring this path length).

To test these hypotheses, we used a dataset (Martínez-García *et al*. 2015) of 10 plant-AMF networks that were sampled at different sites along a long-term (>120,000 yr) chronosequence comprising all stages of ecosystem development, including retrogression (Peltzer *et al*. 2010). Chronosequences represent a powerful tool for understanding long-term co-ordinated changes amongst species and their interactions, resource availability and ecosystem processes (Peltzer *et al*. 2010). Strong gradients from initial N-limitation to eventual strong P-limitation of ecosystem processes drive turnover of plant and AMF species (Richardson *et al*. 2004; Martínez-García *et al*. 2015). We treated ecosystem progression (during which plant species richness and biomass increase; Richardson *et al*. 2004) as the assembly phase, and ecosystem retrogression (during which strong P-limitation drives declines in plant diversity and biomass; Richardson *et al*. 2004) as the network disassembly phase. We defined attachment as the first ecosystem stage at which a plant or AMF species was observed along the chronosequence, and detachment as the final ecosystem stage in which a species was observed.

## Materials and Methods

### Dataset

We generated networks of interactions between vascular plants and arbuscular mycorrhizal fungi (AMF) using data from a study that examined AMF beta diversity and the importance of soil age vs. plant host for explaining AMF community structure (Martínez-García *et al*. 2015). Sampling and sequencing methods can be found in that study or in the Supporting Information (Appendix 1), but we summarise the key points here. Sampling was conducted along the Franz Josef soil chronosequence, on the southern west coast of the South Island, New Zealand. An important feature of this chronosequence, which led us to select this site, is that strong soil nutrient gradients (Appendix 1 in the Supporting Information) are associated with pronounced shifts in ecological community composition, structure and function, such that ecosystem development exhibits a clear progressive phase up to the surface soils aged 5,000 - 12,000 years, then retrogression in the older sites. We treated the sites up to 12,000 years old as the assembly phase, because this was the site of peak tree biomass and height, and a retrogressive phase thereafter in which tree basal area (biomass) declines about three-fold (Richardson *et al*. 2004; Wardle *et al*. 2008) (the disassembly phase). We defined assembly and disassembly phases a priori according to well-characterized ecosystem stages of progression and retrogression (Richardson *et al*. 2004; Wardle *et al*. 2008), rather than by using the size of the sampled plant-AMF network, for two reasons. First, younger sites have many herbaceous plants of low biomass (so many can fit into a given sample area), which can make sampled species richness per plot appear lower in later sites (see Table S1) because the roots of large-biomass trees are more likely to be repeatedly sampled at random. Second, many of the diverse species that were sampled at young sites were not important components of our plant-AMF network, which is evidenced by the sampled plant-fungal associations showing a significant pattern of accumulation up to the site of peak biomass then subtraction during retrogression (described below).

We sampled fungal communities on roots at ten surfaces of the following ages (in years): <5, 15, 70, 290, 500, 1000, 5000, 12,000, 60,000, 120,000 (Supporting Information: Appendix 1 and Table S1). At each site we sampled 50 root fragments, which then underwent a molecular analysis to identify the plant species, as well as any AMF present inside the root (see Martínez-García *et al*. 2015 or Appendix 1 in the Supporting Information). We did not assign weights to these links, as the hypotheses we test here relate to the initial formation of interactions, rather than their strength or frequency once formed.

### Data analysis

We hypothesized that species entering the network for the first time during the progressive phase would associate non-randomly with other existing species based on their direct or indirect connections to other species within the network (i.e. preferential attachment; Barabási & Albert 1999). We also hypothesized that the same would occur during the disassembly phase, such that the probability of species going locally extinct from the network would depend on the network position of species with which they associate (i.e. preferential detachment (Salathé *et al*. 2005; Aizen *et al*. 2012)). We tested for preferential formation of interactions (attachment) by new species during ecosystem progression (5 - 12,000 year-old) and preferential disappearance (detachment) during ecosystem retrogression (12,000 – 120,000 year-old), representing respectively the assembly and disassembly of the network. A species was deemed to have attached during the progressive phase when it first appeared in our samples of the chronosequence. During the progressive phase, once a species had attached (i.e. a link was formed), it was deemed to remain part of the network for the remainder of ecosystem progression. This assumption was made to ensure that any species that appeared, disappeared then reappeared during the progressive phase were not counted twice, because this would give any species-specific interaction preferences of these species a disproportionate weighting in analyses. Also, it gave the largest sample size with which to estimate each species’ number of interaction partners (our predictor variable) in each stage.

Detachment was deemed to have occurred during the retrogressive phase when a species that had been present during the peak of the progressive phase was absent from a given site and all remaining sites along the sequence. Note that, because by definition attachment and detachment processes must occur during the interval between two chronosequence stages, we treated the 12,000 year-old site (which represented the community peak in diversity and biomass) as the ‘peak’ community, comprising the last stage of assembly and the first stage from which disassembly occurred. Although there were only two sites older than this ‘peak’ community, the statistical power for the test came from the number of species within each site, because preferential detachment was tested across species, within sites. An underpinning assumption of this analysis (and of our selection of the 12,000 year site as the peak) is that interactions have the tendency to accumulate during the assembly phase then be lost during the disassembly phase. In other words, the interactions present in sites preceding or following the peak community would be nested subsets of those in the peak site. This pattern has been seen previously during the disassembly of pollination networks (Aizen *et al*. 2012). To test for such a pattern, we generated two matrices in which we ranked the sites according to their age (rows), either becoming progressively older or younger than the site of peak biomass, which was in the top row. Cells of the matrices depicted the presence or absence of each pairwise plant-AMF association (columns). These matrices included all interactions, not just those present in the peak-biomass site. We tested each matrix for nestedness, using the nestedness metric based on overlap and decreasing fill (NODF), with 999 permutations and a swap algorithm, using the oecosimu function in the vegan (Oksanen *et al*. 2008) package for R (R Core Team 2013). This analysis revealed a pattern whereby the peak biomass site had a set of interactions, of which progressively older or younger sites tended to have decreasing subsets (P < 0.001 for rows, columns and overall NODF statistic for both assembly and disassembly), such that interactions, on average, tended to progressively accumulate during progression but be sequentially lost during retrogression.

#### Preferential attachment and detachment

We analyzed preferential attachment and detachment processes using generalized linear mixed effects models with binomial errors and the canonical logit link function, conducted in the lme4 package (Bates *et al*. 2014) in R. We used separate models to test for attachment processes during progression vs. detachment processes during retrogression, and for plants vs. AMF. The response variable in each model was binary, whereby each existing species in the network for each site was coded with a value of one if a newly-arriving species attached to it, or detached from it, in that site, otherwise zero. Site (surface along the chronosequence) was included as a random factor, such that preferential attachment was tested according to the relative network roles of the various species within a site, rather than comparing species across sites, which could be influenced by differences in the sizes of networks. Identity of the attaching species, nested within site, was also included as a random factor to control for the non-independence of a single plant or AMF-OTU attaching or detaching from multiple partners. The fixed predictor variable was either degree or centrality (defined below).

To measure the path length among species of a trophic level, which is related to the number of species through which positive indirect effects (Bastolla *et al*. 2009) must travel, we used closeness centrality (hereafter ‘centrality’, defined in Fig. S1) in the unipartite projection of the bipartite interaction network (Gómez & Perfectti 2012). Centrality is often correlated with degree (Martín González *et al*. 2010) (see also Fig. S3), but carries additional information by measuring the number of links from a focal species to all others in the network (not just immediate neighbors), and has been used as a measure of species’ importance (Martín González *et al*. 2010) or functional specialization (Dalsgaard *et al*. 2008) in mutualistic networks. In the unipartite projection, species at a given trophic level are linked if they share an interaction partner (Fig. S1), so high centrality indicates that a species shares partners with many other species that also share partners with many others. In this sense, high centrality in the unipartite projection (Fig. S1) indicates that positive indirect effects of sharing mutualists with other species at the same trophic level (Bastolla *et al*. 2009) would have a short distance to travel. In contrast, species with low centrality tend to interact with partners that are not used by the dominant consortium of tightly-interacting plants and AMF. Although centrality may be implicitly used to infer the nature of flows through a network (e.g. paths vs. walks, replication vs. spread), we do not make any assumption about the nature of flows, if any, in the plant-AMF network. Rather, we use centrality simply as a measure of the length of indirect paths within a trophic level. Centrality was calculated in the ‘sna’ package (Butts 2014) in R, scaled within a given network (i.e. within each site).

We tested whether attachment to a node during assembly, or detachment from a node during disassembly, depended on that node’s centrality (Fig. S1) or degree (the number of partners with which the species interacts, normalized within networks of a given site age). We used the centrality/degree of a node in the chronosequence stage immediately prior to that in which the new node attached or an existing node detached. This allowed us to compare the fit of models with centrality vs. degree as the predictor using the Akaike Information Criterion (AIC). Thus, the predictor variable in each model was either the degree or centrality of each existing species in the site, to which newly-arriving species either attached vs. did not, or detached vs. did not in the next stage of the chronosequence. For completeness, we also ran models with both of these predictors, to determine if any potential collinearity qualitatively altered their separate effects. None of our models showed any signs of overdispersion. We repeated these analyses controlling for species abundances in case sampling effort generated any spurious effects (see Controlling for the effect of species abundances in Appendix 1 of the Supporting Information).

We omitted from this analysis any links representing new species attaching to other newly-arriving species, because we could not ascribe a value of network position for those species to use as the predictor variable. However, these pairings among two newly-arriving species were rare, and the species involved typically also interacted with other species already present in the network. Across all sites, 20 newly-arriving plant species were associated with newly-arriving AMF, and of these, only 4 did not also attach to AMF species that were already present in the network, for which network position could be calculated. Similarly, of the 15 newly-arriving AMF species that attached to newly-arriving plants, 7 did not also attach to existing species.

### Simulations

To explore the consequences of preferential attachment and detachment based on centrality vs. degree, we conducted a set of simulations of network assembly and disassembly scenarios (more detail provided in Appendix 1 of the Supporting Information). At each time step, newly-arriving plant or AMF species would interact with one existing species, with a probability proportional to the existing species’ degree (i.e. the Barabási & Albert (1999) model) or distance from other species at the same trophic level (closeness centrality). We also ran a third scenario whereby attachment was random (equally probable attachment to any existing species), to provide a null point of comparison.

This assembly phase continued until networks contained 50 species of plant and 50 of AMF. Subsequently, each network was subjected to a disassembly phase, using the same scenario (degree-based, centrality-based or random) as was used during that network’s assembly. Alternating plant and AMF species were removed (i.e. went extinct) with a probability that was inversely proportional to the degree or closeness centrality of the species with which they interacted, such that species that interacted with species of high degree or high centrality were less likely to go extinct. As with the assembly phase, we ran a scenario of the disassembly phase whereby extinction was random.

Each scenario of the assembly and disassembly phases was run for 1000 replicates (each comprising multiple time steps). At each time step within each replicate, after species arrived or went extinct, we recorded the nestedness of the network using NODF, calculated using the nestednodf function in the vegan package (Oksanen *et al*. 2008) for R. Nestedness is a common feature of mutualistic networks (Bascompte *et al*. 2003), which has also been observed recently in plant-mycorrhizal networks (Montesinos-Navarro *et al*. 2012) and has been shown to increase the persistence of networks (Bastolla *et al*. 2009; Thebault & Fontaine 2010). We scaled the observed NODF by the distribution of NODF values obtained by randomizing the adjacency matrix using a null model (Appendix 1 of the Supporting Information).

## Results

In total our samples yielded 33 operational taxonomic units (OTUs, hereafter referred to as species) of AMF and 53 species of plant, among which we observed 399 pairwise interactions (i.e. links, defined as the colonisation of a plant root by an AMF species) (Fig. 1). There were an average of 16 AMF (SD = 3.33) and 10.8 (SD = 3.77) plant species, with 137.5 (SD = 24.14) interactions per site (Table S1). The AMF community changes and turnover along the chronosequence have been described in detail elsewhere (Martínez-García *et al*. 2015).

**Figure 1:**
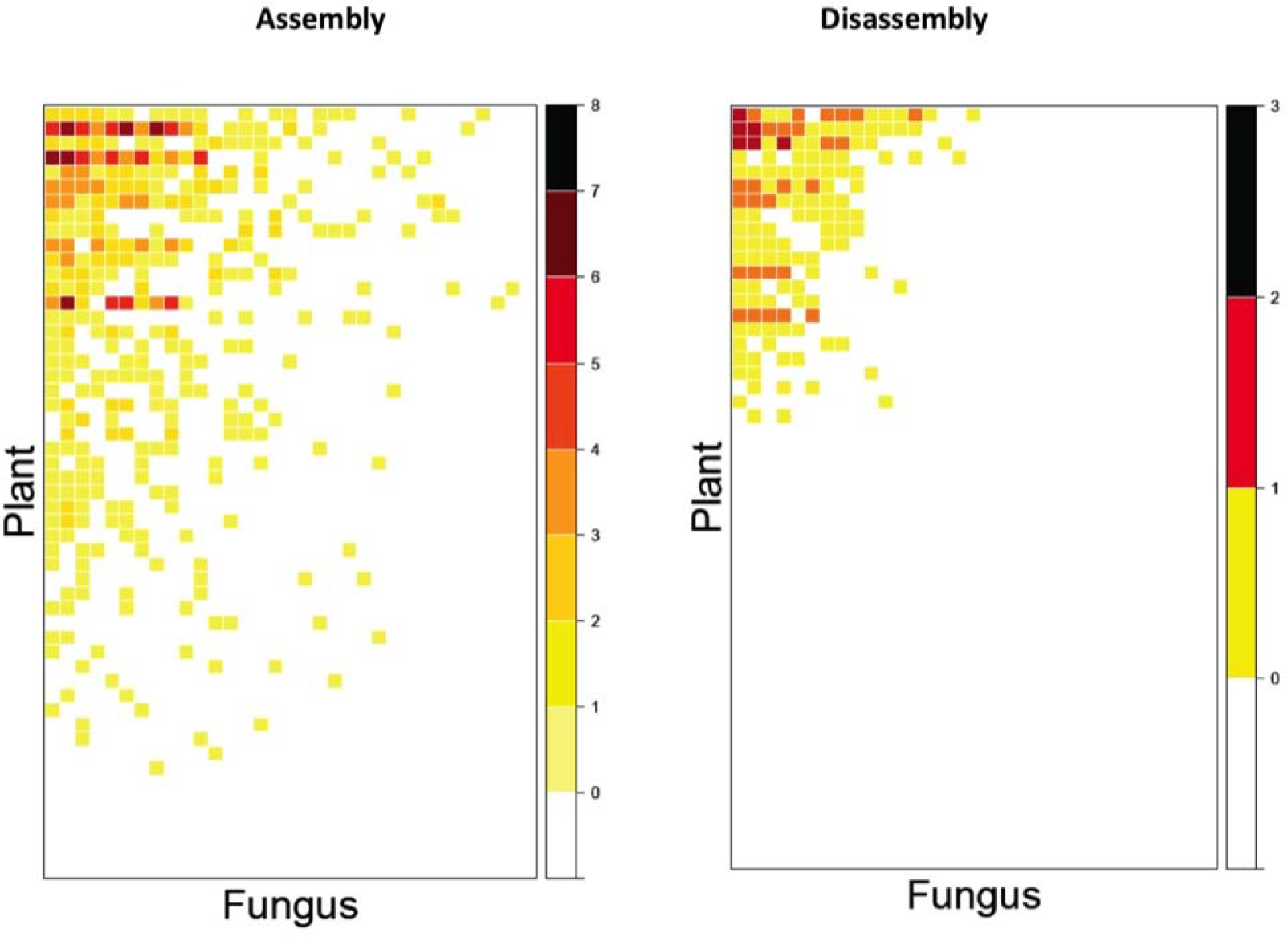
Plant-mycorrhizal association matrices during assembly (N = 8 sites) and disassembly (i.e. retrogressive, N = 3 sites) phases of ecosystem development. The site of peak biomass represents the end of assembly and beginning of disassembly, so is included in both figures. Darker colours indicate that the plant-mycorrhizal association was present in a greater number of sites (i.e. it formed early during assembly or persisted during retrogression). These darker interactions tend to involve species that share interaction partners with many others that also share partners with each other (i.e. species with high centrality). The network is significantly nested (see Appendix 2 of the Supporting Information), whereby specialists interact with proper subsets of the species that interact with generalists.

We first tested whether preferential attachment or detachment occurred to existing species that had many partners or high centrality. At their earliest occurrence along the sequence, both plant and AMF species were significantly more likely to interact with species that shared interaction partners with many others (i.e. high centrality, Fig. 2A, P < 0.0001 in both cases), but not those with many partners (i.e. high degree; P > 0.862 in both cases) (Table S2). In addition to this preferential attachment, plants also showed preferential detachment, such that they were significantly less likely to be lost from the network during ecosystem retrogression if they interacted with AMF species that had high centrality (P < 0.0001; Fig. 2D; Table S2). In contrast, the apparent decline in AMF detachment probability with plant centrality (Fig. 2C) or degree was non-significant (Table S2). Thus, symmetrical assembly and disassembly processes, either of which could theoretically generate complex network architecture (Barabási & Albert 1999; Salathé *et al*. 2005; Bastolla *et al*. 2009; König *et al*. 2012), both operated on the plant community to rapidly generate architecture that did not change significantly throughout ecosystem development (Appendix 2 of the Supporting Information, Table S3, Fig. S2).

**Figure 2:**
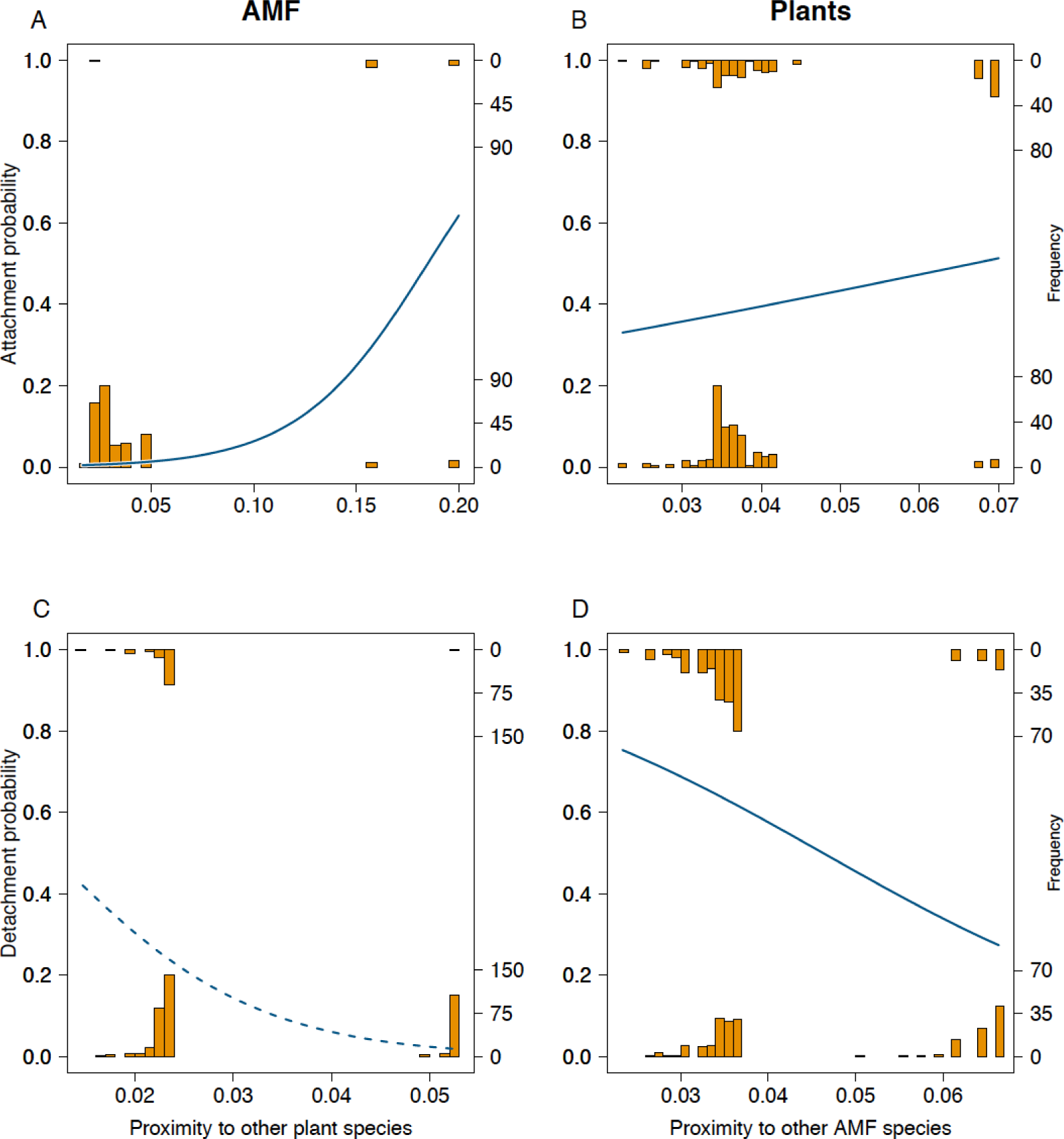
Probability (left vertical axis) that a new A,C) arbuscular mycorrhizal fungal (AMF), or B,D) plant species entering the network will interact with an existing species during assembly (top), or detach from a species during disassembly (bottom) as a function of that species’ proximity in the network to others of its trophic level (measured as closeness centrality of the unipartite projection of the plant-AMF network, see Fig. S1). Histograms top and bottom of each graph represent the frequencies (right vertical axis) of ones and zeroes respectively in the raw data. Trend lines are based on inverse-linked coefficients of a binomial linear mixed effects model with chronosequence stage (plant-mycorrhizal network) as a random effect. Solid lines were statistically significant (P < 0.025 in all cases), dashed line was not significant at alpha = 0.05.

Although species with many direct interaction partners also had significantly shorter indirect paths on average (i.e. degree and centrality were correlated in our networks; Fig. S3), we found no evidence of preferential attachment or detachment of plants or AMF based on degree alone (Table S2). Moreover, when both degree and centrality were included in models, attachment probability *decreased* with increasing degree (Table S2) and plant detachment probability *increased* when they interacted with AMF of high degree (Table S2), as has been observed in a long-term study of pollination networks (Burkle *et al*. 2013). Importantly, in all our models, centrality explained attachment or detachment probability better than did equivalent models with only degree as the predictor, and its effect remained qualitatively unchanged when degree was also included in the model (Table S2). Most importantly, the relation between AMF centrality and plant attachment and detachment probability remained significant after controlling for abundance (Appendix 2 of the Supporting Information, Table S4).

Our findings suggest that both attachment and detachment are key processes in network development and, when combined, the loss or gain of links becomes self-reinforcing (König *et al*. 2012). We simulated the processes examined here to explore the consequences for network architecture (Appendix 1 of the Supporting Information) and found that both preferential attachment and detachment processes were important for determining and maintaining the nestedness of the network (Fig. S4), a key element of ecological network architecture (Bastolla *et al*. 2009; Montesinos-Navarro *et al*. 2012) whereby specialists interact with species that also interact with generalists. Preferential attachment (based on degree or centrality) generated networks that were considerably more nested than random, and preferential detachment maintained this nestedness in the face of species extinctions (Fig. S4).

## Discussion

Obviously species do not have information about the architecture of their interaction networks, so centrality (a species’ indirect proximity to others at the same trophic level) must be associated with ecological characteristics that make species more likely to attract and retain interaction partners. There are several potential, non-exclusive explanations for this attractiveness. The simplest would be if abundant species shared many partners due to a high probability of random encounters. In our dataset, there was a weak correlation between a species’ abundance and centrality within sites (Fig. S5). Moreover, the core of interacting generalists (i.e. the species involved in the majority of interactions in the network, and whose interaction partners overlapped considerably with the rest of the community) was present throughout much of ecosystem development (see dark colors in top left of matrices in Fig. 1), and this ubiquity likely made them a reliable target for newly arriving species. However, causality cannot be inferred from these correlations in our dataset, as attraction of diverse fungal mutualists could equally cause longer-term persistence of plants (Scheublin *et al*. 2007). Moreover, the significant relationships between centrality and attachment or detachment probability remained qualitatively unchanged when controlling for abundance (Appendix 2 of the Supporting Information, Table S4).

Therefore, although abundance likely plays some role, the attractiveness of plant species with high centrality to AMF, and vice versa, reflects additional benefits to associating with those species. One such benefit of close proximity is the potential to receive beneficial indirect effects (Bastolla *et al*. 2009); plants may benefit from the AMF communities generated by other species (Bever 2002), and plant-plant competition can be reduced (and plant biomass increased) by high AMF diversity (Wagg *et al*. 2011). Thus, consortia of plants that are able to share diverse AMF communities with one another may grow better, and therefore have the ability to provide more carbon to AMF symbionts. These groupings may also reflect habitat preference of the species involved, with habitat generalist plant and AMF species interacting tightly during early succession and forest specialists during late succession (Davison *et al*. 2011), an hypothesis congruent with findings of partner specificity occurring at the level of ecological groups (Öpik *et al*. 2009).

Plant species were also less likely to be lost during retrogression if they interacted with AMF species that had high centrality (Fig. 2D). We hypothesized (based on the benefits described above) that mutually beneficial interactions will involve species with high centrality, and we observed that plant species that preferentially interacted with these core species were significantly more likely to persist during retrogression. The decline in soil P during ecosystem development likely makes plants increasingly dependent on AMF-provided P. Likewise, declining plant photosynthetic rates along the chronosequence (Whitehead *et al*. 2005) could increase competition among AMF for plant-derived carbon. Thus, the need to develop and maintain mutualists and avoid symbiotic ‘cheaters’ will intensify along the sequence, particularly during retrogression. Over time, both plants and AMF can reduce their resource allocation to less-beneficial partners (Kiers *et al*. 2011), particularly when resources are scarce (Bever 2015). The interactions that persist during retrogression may therefore be those that are most mutually beneficial. Previous work has found that species in late-successional plant-AMF networks tend to interact more frequently with a subset of their total range of partners (Bennett *et al*. 2013), and a viable strategy may thus be to test a number of partners initially, then gradually restrict resource allocation to a subset of these that provide the greatest benefit (Bever 2015) as resources become limiting. This process could easily occur at the level of plant individuals, but be reflected as a loss of links at the species level.

Finally, if our findings are found to generalize to other ecological networks, they would have several potential applications. First the symmetry of assembly and disassembly processes for plants and our observation that interactions in sites moving forward or backwards from the site of peak biomass were nested subsets of that site (see Materials and Methods section) suggests that knowledge of assembly order for a given system could be used to predict the interactions most at risk of extinction, such as in using network information for conservation (Tylianakis *et al*. 2010). Species reintroductions during restoration could also follow the reverse order of extinction and replace extinct species with others that fulfill the same network role, or focus on species that share interaction partners with others, and will then attract interactions with any new species that colonize. Similarly, non-native species should interact preferentially with high-partner-overlap species in the network, which could therefore be a focus for biosecurity monitoring to detect invasions early.

It is impossible to fully understand the architecture of complex networks in isolation from the dynamic processes that generate and maintain them (Barabási & Albert 1999). We have shown empirically that both preferential attachment and detachment underpinned the development of an ecological network under changing abiotic conditions. Importantly, species with short indirect paths to others at the same trophic level were more attractive interaction partners, congruent with suggestions that positive indirect effects may be important for species persistence in mutualistic networks (Bastolla *et al*. 2009). Moreover, interactions in sites moving forward or backward in age from the site of peak biomass were nested subsets of those in the peak biomass site. This pattern has been observed in the disassembly of pollination networks (Aizen *et al*. 2012), and we found it to also occur during assembly. Finally, we showed through simulation that symmetrical, self-reinforcing processes can generate and maintain important features of interaction network architecture (Barabási & Albert 1999; Salathé *et al*. 2005; Bastolla *et al*. 2009; König *et al*. 2012), thereby linking species colonization and extinction processes to emergent and potentially stabilizing (Bastolla *et al*. 2009; Thebault & Fontaine 2010) ecosystem properties.

## Acknowledgements

We thank P.J. Bellingham, T. Fukami, J.H. Jones, S. Pawar, D.B. Stouffer and members of the Tylianakis/Stouffer lab for critical discussions and comments on the manuscript. N. Bolstridge and C. Mitchel provided valuable lab assistance. The research was funded by Core funding for Crown Research Institutes from the New Zealand Ministry of Business, Innovation and Employment's Science and Innovation Group, a Rutherford Discovery Fellowship to JMT, and the Bio-protection Research Centre (JMT and IAD).

## Author information

The authors declare no competing financial interests. Correspondence and requests for materials should be addressed to J.M.T.

**Supplementary Information** is available online

## References

Aizen, M.A., Sabatino, M. & Tylianakis, J.M. (2012). Specialization and Rarity Predict Nonrandom Loss of Interactions from Mutualist Networks. Science, 335, 1486–1489.

Barabási, A.-L. & Albert, R. (1999). Emergence of scaling in random networks. Science, 286, 509–512.

Bascompte, J., Jordano, P., Melian, C.J. & Olesen, J.M. (2003). The nested assembly of plant-animal mutualistic networks. Proceedings of the National Academy of Sciences of the United States of America, 100, 9383–9387.

Bascompte, J. & Stouffer, D.B. (2009). The assembly and disassembly of ecological networks. Philosophical Transactions of the Royal Society B-Biological Sciences, 364, 1781–1787.

Bastolla, U., Fortuna, M.A., Pascual-Garcia, A., Ferrera, A., Luque, B. & Bascompte, J. (2009). The architecture of mutualistic networks minimizes competition and increases biodiversity. Nature, 458, 1018–U1091.

Bates, D., Maechler, M., Bolker, B. & Walker, S. (2014). lme4: Linear mixed-effects models using Eigen and S4.

Bennett, A.E., Daniell, T.J., Öpik, M., Davison, J., Moora, M., Zobel, M. et al. (2013). Arbuscular mycorrhizal fungal networks vary throughout the growing season and between successional stages. PloS one, 8, e83241.

Bever, J.D. (2002). Negative feedback within a mutualism: host–specific growth of mycorrhizal fungi reduces plant benefit. Proceedings of the Royal Society of London B: Biological Sciences, 269, 2595–2601.

Bever, J.D. (2015). Preferential allocation, physio‐evolutionary feedbacks, and the stability and environmental patterns of mutualism between plants and their root symbionts. New Phytologist, 205, 1503–1514.

Brundrett, M.C. (2009). Mycorrhizal associations and other means of nutrition of vascular plants: understanding the global diversity of host plants by resolving conflicting information and developing reliable means of diagnosis. Plant and Soil, 320, 37–77.

Burkle, L.A., Marlin, J.C. & Knight, T.M. (2013). Plant-pollinator interactions over 120 years: loss of species, co-occurrence, and function. Science, 339, 1611–1615.

Butts, C.T. (2014). sna: Tools for Social Network Analysis.

Dalsgaard, B., Martín González, A.M., Olesen, J.M., Timmermann, A., Andersen, L.H. & Ollerton, J. (2008). Pollination networks and functional specialization: a test using Lesser Antillean plant–hummingbird assemblages. Oikos, 117, 789–793.

Davison, J., Öpik, M., Daniell, T.J., Moora, M. & Zobel, M. (2011). Arbuscular mycorrhizal fungal communities in plant roots are not random assemblages. FEMS Microbiology Ecology, 78, 103–115.

Díaz-Castelazo, C., Guimarães, P.R., Jordano, P., Thompson, J.N., Marquis, R.J. & Rico-Gray, V. (2010). Changes of a mutualistic network over time: reanalysis over a 10‐year period. Ecology, 91, 793–801.

Ehinger, M., Koch, A.M. & Sanders, I.R. (2009). Changes in arbuscular mycorrhizal fungal phenotypes and genotypes in response to plant species identity and phosphorus concentration. New Phytologist, 184, 412–423.

Encinas-Viso, F., Alonso, D., Klironomos, J.N., Etienne, R.S. & Chang, E.R. (2015). Plant–mycorrhizal fungus co‐occurrence network lacks substantial structure. Oikos.

Gómez, J.M. & Perfectti, F. (2012). Fitness consequences of centrality in mutualistic individual-based networks. Proceedings of the Royal Society B: Biological Sciences, 279, 1754–1760.

Gómez, J.M., Verdú, M. & Perfectti, F. (2010). Ecological interactions are evolutionarily conserved across the entire tree of life. Nature, 465, 918–921.

Horn, S., Caruso, T., Verbruggen, E., Rillig, M.C. & Hempel, S. (2014). Arbuscular mycorrhizal fungal communities are phylogenetically clustered at small scales. The ISME journal, 8, 2231–2242.

Jordano, P., Bascompte, J. & Olesen, J.M. (2003). Invariant properties in coevolutionary networks of plant-animal interactions. Ecology letters, 6, 69–81.

Kiers, E.T., Duhamel, M., Beesetty, Y., Mensah, J.A., Franken, O., Verbruggen, E. et al. (2011). Reciprocal rewards stabilize cooperation in the mycorrhizal symbiosis. science, 333, 880–882.

König, M., Tessone, C. & Zenou, Y. (2012). Nestedness in Networks: A Theoretical Model and Someapplications. SIEPR Discussion Papers, 11, 1–61.

Krüger, M., Teste, F.P., Laliberté, E., Lambers, H., Coghlan, M., Zemunik, G. et al. (2015). The rise and fall of arbuscular mycorrhizal fungal diversity during ecosystem retrogression. Mol. Ecol., 24, 4912–4930.

Martín González, A.M., Dalsgaard, B. & Olesen, J.M. (2010). Centrality measures and the importance of generalist species in pollination networks. Ecological Complexity, 7, 36–43.

Martínez-García, L.B., Richardson, S.J., Tylianakis, J.M., Peltzer, D.A. & Dickie, I.A. (2015). Host identity is a dominant driver of mycorrhizal fungal community composition during ecosystem development. New Phytologist, 205, 1565–1576.

Moeller, H.V. & Neubert, M.G. (2016). Multiple friends with benefits: an optimal mutualist management strategy? The American Naturalist, 187, E1–E12.

Montesinos-Navarro, A., Segarra-Moragues, J.G., Valiente-Banuet, A. & Verdu, M. (2012). The network structure of plant-arbuscular mycorrhizal fungi. New Phytologist, 194, 536–547.

Newman, E.I. (1988). Mycorrhizal links between plants: their functioning and ecological significance. Advances in Ecological Research, 18, 243–270.

Oksanen, J., Kindt, R., Legendre, P., O'Hara, B., Simpson, G.L., Solymos, P. et al. (2008). vegan: Community Ecology Package http://cran.r-project.org/.

Olesen, J.M., Bascompte, J., Elberling, H. & Jordano, P. (2008). Temporal dynamics in a pollination network. Ecology, 89, 1573–1582.

Öpik, M., Metsis, M., Daniell, T., Zobel, M. & Moora, M. (2009). Large‐scale parallel 454 sequencing reveals host ecological group specificity of arbuscular mycorrhizal fungi in a boreonemoral forest. New Phytologist, 184, 424–437.

Peltzer, D.A., Wardle, D.A., Allison, V.J., Baisden, W.T., Bardgett, R.D., Chadwick, O.A. et al. (2010). Understanding ecosystem retrogression. Ecological Monographs, 80, 509–529.

Ponisio, L.C., Gaiarsa, M.P. & Kremen, C. (2017). Opportunistic attachment assembles plant–pollinator networks. Ecology Letters, 20, 1261–1272.

Powell, J.R., Karunaratne, S., Campbell, C.D., Yao, H., Robinson, L. & Singh, B.K. (2015). Deterministic processes vary during community assembly for ecologically dissimilar taxa. Nature communications, 6.

R Core Team (2013). R: A language and environment for statistical computing. R Foundation for Statistical Computing, Vienna Austria.

Richardson, S.J., Peltzer, D.A., Allen, R.B., McGlone, M.S. & Parfitt, R.L. (2004). Rapid development of phosphorus limitation in temperate rainforest along the Franz Josef soil chronosequence. Oecologia, 139, 267–276.

Salathé, M., May, R.M. & Bonhoeffer, S. (2005). The evolution of network topology by selective removal. Journal of the Royal Society Interface, 2, 533–536.

Scheublin, T.R., Van Logtestijn, R.S.P. & Van Der Heijden, M.G.A. (2007). Presence and identity of arbuscular mycorrhizal fungi influence competitive interactions between plant species. Journal of Ecology, 95, 631–638.

Simard, S.W. & Durall, D.M. (2004). Mycorrhizal networks: a review of their extent, function, and importance. Canadian Journal of Botany, 82, 1140–1165.

Suding, K.N. & Hobbs, R.J. (2009). Threshold models in restoration and conservation: a developing framework. Trends in ecology & evolution, 24, 271–279.

Thebault, E. & Fontaine, C. (2010). Stability of Ecological Communities and the Architecture of Mutualistic and Trophic Networks. Science, 329, 853–856.

Thompson, R.M., Brose, U., Dunne, J.A., Hall Jr, R.O., Hladyz, S., Kitching, R.L. et al. (2012). Food webs: reconciling the structure and function of biodiversity. Trends in ecology & evolution.

Tylianakis, J., Laliberté, E., Nielsen, A. & Bascompte, J. (2010). Conservation of species interaction networks. Biological Conservation, 143, 2270–2279.

Vályi, K., Rillig, M.C. & Hempel, S. (2015). Land - use intensity and host plant identity interactively shape communities of arbuscular mycorrhizal fungi in roots of grassland plants. New Phytologist, 205, 1577–1586.

Wagg, C., Jansa, J., Stadler, M., Schmid, B. & Van Der Heijden, M.G.A. (2011). Mycorrhizal fungal identity and diversity relaxes plant-plant competition. Ecology, 92, 1303–1313.

Wardle, D., Bardgett, R., Walker, L., Peltzer, D. & Lagerström, A. (2008). The response of plant diversity to ecosystem retrogression: evidence from contrasting long-term chronosequences. Oikos, 117, 93–103.

Whitehead, D., Boelman, N.T., Turnbull, M.H., Griffin, K.L., Tissue, D.T., Barbour, M.M. et al. (2005). Photosynthesis and reflectance indices for rainforest species in ecosystems undergoing progression and retrogression along a soil fertility chronosequence in New Zealand. Oecologia, 144, 233–244.

